# The tumor suppressor FBW7 and the vitamin D receptor are mutual cofactors

**DOI:** 10.1101/272500

**Authors:** Reyhaneh Salehi-Tabar, Babak Memari, Hilary Wong, Natacha Rochel, John H. White

**Affiliations:** Departments of Physiology, McGill University, Montreal Qc, Canada.; Departments of Medicine, McGill University, Montreal Qc, Canada.; Department of Integrative and Structural Biology, Institut de Génétique et de Biologie Moléculaire et Cellulaire, Illkirch, France.

**Keywords:** Cell cycle regulation, E3 ligase, Protein turnover, Transcription regulation, VDR

## Abstract

The E3 ligase FBW7 targets drivers of cell cycle progression such as c-MYC for proteasomal degradation. It is frequently mutated in cancer, and is a tumor suppressor. Extensive epidemiological data links vitamin D deficiency to increased incidence of several cancers, although the underlying cancer-preventive mechanisms are poorly understood. Here, we show that hormonal 1,25-dihydroxyvitamin D3 (1,25D) rapidly stimulates the interaction of the VDR with FBW7, and that of FBW7 with c-MYC. In contrast, it blocks the association of FBW7 with c-MYC antagonist MXD1. 1,25D also enhances the association of FBW7, proteasome subunits, and ubiquitin with DNA-bound c-MYC, consistent with induced degradation of c-MYC on DNA. In addition to c-MYC, 1,25D accelerates the turnover of other FBW7 target proteins. Intriguingly, FBW7 is essential for optimal *VDR* gene expression. It is also recruited to VDR targets genes, and its depletion attenuates 1,25D-stimulated VDR DNA binding, transactivation, and cell cycle arrest. Thus, the VDR and FBW7 are mutual cofactors, which provides a molecular basis for the cancer-preventive actions of vitamin D through accelerated turnover of FBW7 target proteins.

## Introduction

It is the essential role of E3 ligases of the ubiquitin proteasome system (UPS) to recognize proteins targeted for ubiquitination. SCFs (Skp1, Cullin-1, F box protein) are a class of E3 ligases in which Cullin-1 is a scaffold and the F-box protein is the substrate recognition subunit. UPS components are targeted frequently by oncogenic mutations in cancer [1]–[3]. FBW7 (FBXW7/Sel-10/Ago/hCDC4) is of particular interest because it targets a network of proteins that drive cell cycle progression for proteasomal degradation. FBW7 is one of the most commonly dysregulated UPS proteins in multiple cancer types and is a tumor suppressor [3]–[5]. It recognizes its substrates through so-called “phospho-degron” motifs (consensus T/S-P-X-X-S/T/E), which are phosphorylated by GSK3β [6], [7]. Multiple FBW7 target proteins are oncogenes implicated in cell cycle progression, including cyclin E, the coactivator AIB1 (Amplified in breast cancer 1; a.k.a. SRC1, ACTR), the transcription factor c-Jun, and the apoptosis regulator MCL1 (Myeloid Cell Leukemia 1) [8], [9]. One of the best characterized FBW7 substrates is the oncogenic transcription factor c-MYC, whose expression is dysregulated in ~50% of all cancers [10]. Down-regulation of FBW7 results in accumulation of cellular and chromatin-bound c-MYC [11]. Thr58 of the c-MYC phospho-degron is one of most frequently mutated residues in lymphoma cells [12]. c-MYC activity can be antagonized by the transcriptional repressor MXD1 [13], which, like c-MYC, binds cognate E-boxes as a heterodimer with MAX [14]. c-MYC is expressed in proliferating cells, whereas MXD1 is found mostly in differentiated cells [13], [15].

We have shown that similar to c-MYC, MXD1 can be targeted for degradation by FBW7 [16]. Moreover, we found that the vitamin D receptor (VDR) can differentially regulate the stability of c-MYC and MXD1, and that this regulation is abrogated by ablation of FBW7 [16]. Numerous studies have linked vitamin D deficiency to increased incidence of cancers [17]–[19], The α-helical ligand-binding domain controls transcriptional regulation by the VDR [20], [21]. Notably, the domain contains a flexible 23-residue “insertion” between helices 2 and 3, which is unique to VDR [22]. Intriguingly, 1,25D enhances c-MYC turnover, and reduces that of MXD1, drastically altering the ratio of the two proteins in treated cells [16]. These data suggest that the regulation of FBW7 may be a key contributor to the anti-cancer properties of vitamin D. Here, we show that the VDR and FBW7 are mutual cofactors. 1,25D can enhance the association of FBW7 with c-MYC and accelerates the turnover of several FBW7 target proteins. In addition, FBW7 is essential for optimal VDR expression. Moreover, it is recruited to VDR target genes in a hormone-dependent fashion, and its ablation compromises VDR transactivation, and the capacity of 1,25D to arrest the proliferation of cancer cells. These results suggest that the association of the VDR with FBW7 is a fundamental component of the anti-cancer properties of 1,25D signaling.

## Results and Discussion

c-MYC is one of the best characterized FBW7 substrates [23], and we found that, unexpectedly, turnover of it antagonist MXD1 is also regulated by FBW7 [16]. Furthermore, we observed that 1,25D enhanced c-MYC turnover in a manner that was dependent on FBW7 expression, but that 1,25D had the opposite effect on MXD1 protein expression, stability, and DNA binding [16]. This suggested that the hormone-bound VDR acted as an FBW7 cofactor by regulating access of FBW7 to its substrates. We investigated VDR-MYC interactions by CoIP, which revealed partially hormone-dependent association of the two proteins (Fig. 1A). Similarly, GST pull-down assays with free GST or GST fused to the full-length VDR showed that the receptor and c-MYC interacted in a partially hormone-dependent manner (Fig. 1B). We also investigated the effect of 1,25D on the association of FBW7 with c-MYC. As commercially available antibodies do not reliably recognize endogenous FBW7, SCC25 cells were lentivirally transduced with an expression vector for Flag-tagged FBW7. Importantly, these experiments revealed that the CoIP of Flag-FBW7 with c-MYC was enhanced by 1,25D (Fig. 1C). Under these conditions, 1,25D treatment induced a modest decline in FBW7 transcripts (Fig. S1A), strongly suggesting that its effect on the association of FBW7 with c-MYC was not due to elevated FBW7 protein expression.

**Figure 1:**
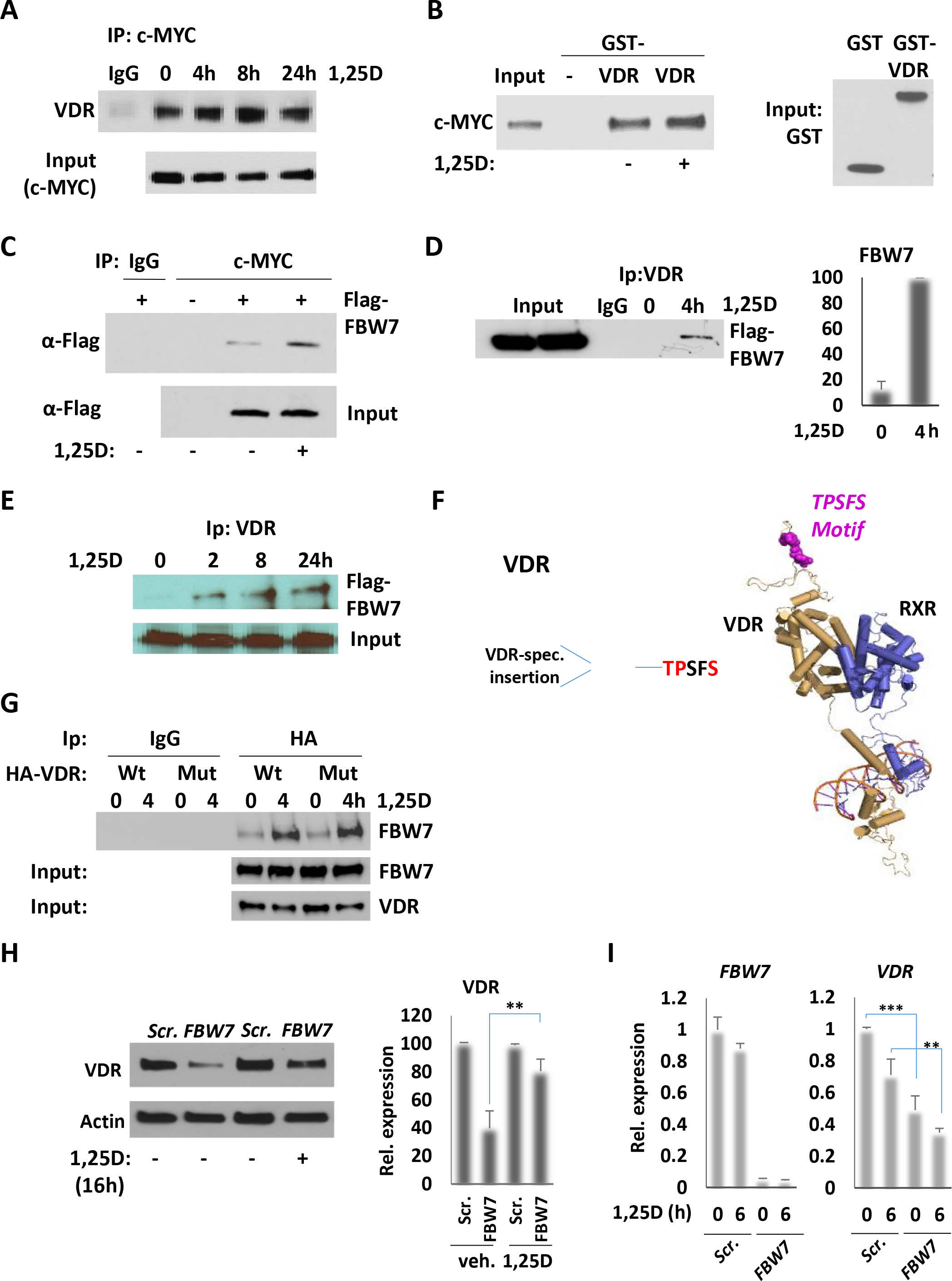
1,25D-dependent Interaction of VDR and MYC with FBW7. **A)** Western analysis of VDR Co-IPed with IgG or c-MYC following treatment of SCC25 cells with 100nM 1,25D. **B)** Left: Western analysis of c-MYC pulled down by GST or GST-VDR −/+1,25D. Right: Western analysis of GST and GST-VDR as input. **C)** Western analysis of Flag-FBW7 Co-IPed with IgG or c-MYC, following treatment of SCC25 cells with 1,25D. Cells were transfected with empty vector or Flag-*FBW7* and treated with 100nM of 1,25D for 6 h. c-MYC was IPed from SCC25 lysate. The total lysate as “input” and IPs were probed for Flag. **D)** Left and **E)** Western analysis of Flag-FBW7 Co-IPed with IgG or VDR, following treatment of SCC25 cells with 100nM of 1,25D, as indicated. The total lysate as “input” and IPs were probed for Flag. **D)** Right: Quantification of Western analysis of Flag-FBW7 Co-IPed with IgG or VDR. **F)** Left: Representation of the VDR and location of the phospho-degron motif TPSFS. Right: Tertiary structure of VDR/RXR showing the phospho-degron in the insertion domain. **G)** Western analysis of Flag-FBW7 Co-IPed with IgG VDR, or mutated VDR, following treatment of SCC25 cells with 100nM of 1,25D for 4 h. Total lysate was probed for Flag or VDR as input. **H)** Left: Western analysis of VDR in FBW7 ablated SCC25 cells. Cells were transfected for *FBW7* or scrambled siRNA and treated with 100nM 1,25D. Actin was used as an internal control. Right: Quantification of Western analysis of VDR. **I)** RT-qPCR assay of *FBW7* and *VDR* mRNA expression after knockdown of *FBW7* in SCC25 cells and treatment with 100nM of 1,25D. **P≤0.01, ***P≤0.001 as determined by One-way ANOVAs followed by Tukey’s post hoc test for multiple comparisons.

Similar studies were performed to determine the effects of 1,25D on the interactions between the VDR, MXD1 and FBW7. Under conditions where MXD1 levels were increasing in the presence of 1,25D (Fig. S1B), its association with the VDR over 8h was largely unchanged but declined after 24h (Fig. S1C). GST pull-down assays revealed that the VDR and MXD1 interacted directly in the absence of hormone, and that 1,25D somewhat reduced VDR-MXD1 binding (Fig. S1D). In the absence of 1,25D, MXD1 expression levels are low [16] (Fig. S1B). However, we were able to detect CoIP of Flag-FBW7 with MXD1 in vehicle-treated cells, which was disrupted by addition of 1,25D (Fig.S1E) under conditions of increasing MXD1 expression. The opposing effects of 1,25D on the association of FBW7 with c-MYC and MXD1 are consistent with the destabilization of c-MYC and stabilization of MXD1 in the presence of hormone.

To probe the interaction of FBW7 with the VDR, we performed CoIPs of SCC25 cells transduced with a lentivector expressing FLAG-FBW7, which revealed a rapid and strongly 1,25D-dependent association of the VDR with FBW7 (Figs. 1D and E). Remarkably, screening of the VDR sequence revealed a consensus FBW7 phospho-degron in an unstructured linker region lying being helices 2 and 3 of the ligand binding domain (LBD; Fig. 1F). This insertion is specific to the VDR among nuclear receptors and is conserved in primates, mice and rats (Fig. S2A). However, CoIPs showed that mutation of the putative phosphodegron had no effect on the 1,25D-dependent interaction of the VDR with FBW7 (Fig. 1G). We further examined the effect of *FBW7* depletion on VDR expression and function. si-RNA knockdown generally led to a near-complete loss of *FBW7* transcripts (e.g. see Fig. 1I). Intriguingly, depletion of FBW7 led to a substantial reduction in VDR protein, an effect that was partially alleviated by addition of 1,25D (Fig. 1H). In the absence or presence of 1,25D, the turnover of the VDR in cycloheximide (CHX)-treated cells was unaffected or modestly accelerated by depletion of FBW7 (Figs. S3A and B), and 1,25D may stabilize the VDR in cells lacking FBW7, although the effect did not reach statistical significance over 24h (Fig. S3C). The observation that phospho-degron mutation did not disrupt VDR-FBW7 interaction, and the lack of increase in VDR stability in FBW7-depleted cells are not consistent with FBW7 recognizing the VDR as a substrate. We then analyzed the effects of FBW7 ablation on *VDR* mRNA levels, which revealed an ~2-fold reduction in transcript levels (Fig. 1I). These results suggest that FBW7 may control the turnover of factor(s) that act to suppress *VDR* mRNA expression, and that diminished *VDR* transcript levels account for much of the loss of VDR protein in absence of FBW7.

Given these findings, we examined the effects of FBW7 ablation on VDR function as a transactivator. Short-term (6h) 1,25D-dependent induction of several direct VDR target genes was compromised in FBW7-depleted cells (Fig. 2A), and ChIP assays revealed that VDR binding to target promoter VDREs after 2h of 1,25D treatment was substantially diminished in the absence of FBW7 (Fig. S4A). Next, we investigated whether FBW7 is recruited to VDREs in the presence of 1,25D. Given the lack of an appropriate antibody to analyze FBW7 by ChIP, we lentivirally transduced SCC25 cells with a vector expressing tagged FBW7. Under these conditions, ChIP assays revealed 1,25D-dependent recruitment of tagged FBW7 to VDR binding sites of target several genes (Fig. 2B). We also tested for association of proteasome subunits SKP1 and RBX1 with VDR target genes by ChIP, which revealed patterns of recruitment similar to those of FBW7 (Figs. 2C and D). Taken together, this suggests that in addition to acting to maintain expression of the VDR, FBW7 functions as a cofactor in VDR-dependent transactivation through recruitment of proteasome subunits to target promoters. The results also infer that 1,25D-dependent gene transcription would be attenuated by proteasome inhibition. As blocking the proteasome may affect the expression/function of several classes of transcription factors, we tested the effects of inhibitors MG132 and Bortezomib on transcription of *CYP24A1* and *CAMP*, whose expression is primarily regulated by the hormone-bound VDR, and found that 1,25D-dependent induction of both genes was compromised (Fig. S5A). Thus, proteasome recruitment and function is an integral part of VDR-dependent transactivation. Finally, to test for possible functional consequences of diminished VDR expression and function in the absence of FBW7, we examined the effect of FBW7 ablation on the anti-proliferative effects of 1,25D in SCC25 cells. These experiments revealed that FBW7 depletion enhanced SCC25 cell proliferation as measured by EdU incorporation and strongly attenuated the capacity of 1,25D to arrest cell proliferation (Fig. 2E and F).

**Figure 2:**
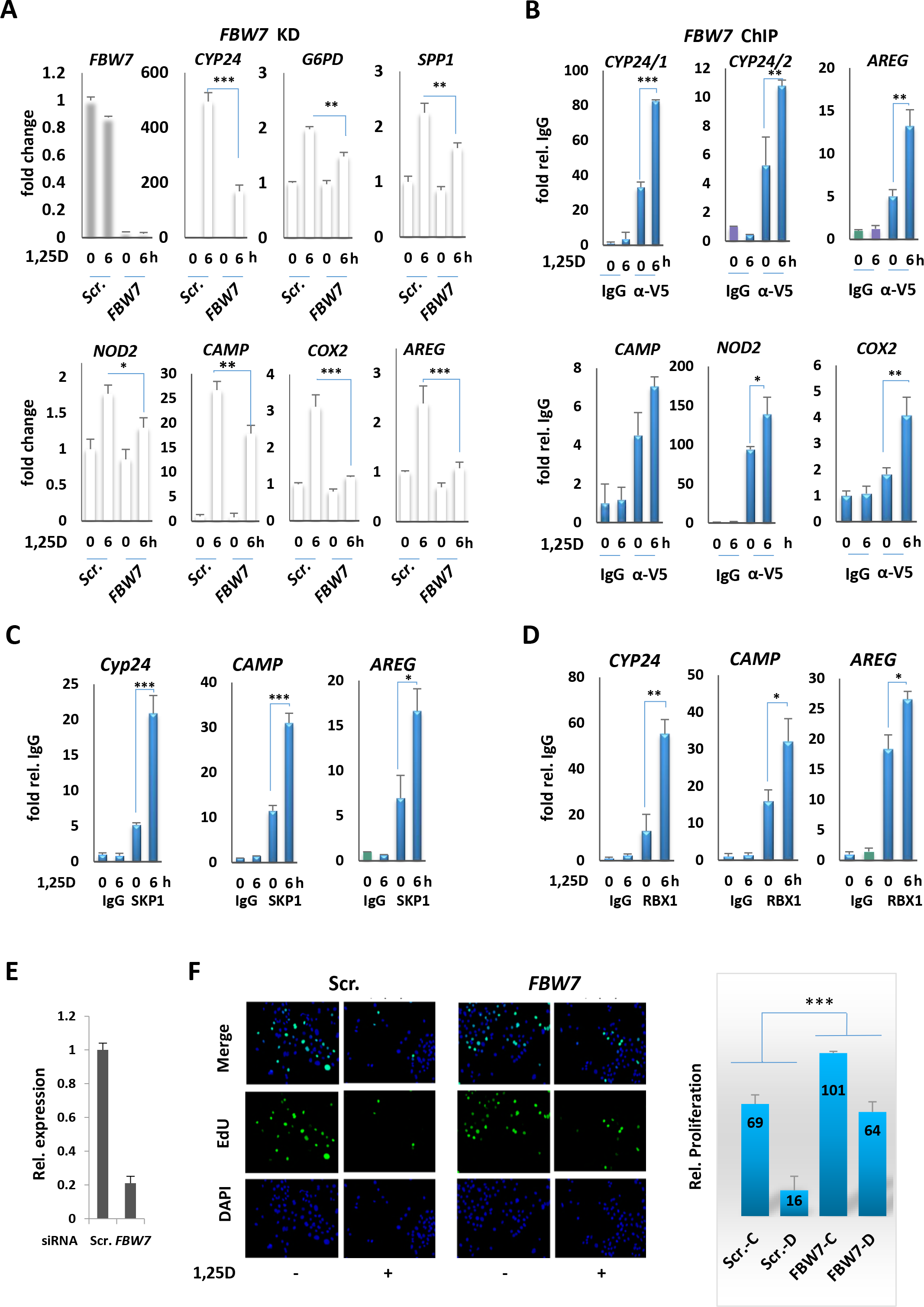
FBW7 as a cofactor on VDR target genes. **A)** RT-qPCR of VDR target gene transcription in SCC25 cells transfected with scrambled or *FBW7* siRNA and treated with 100Nm 1,25D. **B)** Analysis of V5-FBW7 recruitment to VDREs by ChIP-qPCR in V5-FBW7 SCC25 cells treated with 100nM of 1,25D. **C)** and **D)** Analysis of recruitment of SKP1 and RBX1 to VDREs by ChIP-qPCR in SCC25 cells treated with 100nM 1,25D. **E)** RT-qPCR analysis of *FBW7* transcription in SCC25 cells transfected with scrambled or *FBW7* siRNA. **F)** Left: EdU cell proliferation assay in SCC25 cells transfected with scrambled or *FBW7* siRNA and treated with vehicle or 100nM of 1,25D. Right: quantification of EdU cell proliferation assay. *P≤0.05, **P≤0.01, ***P≤0.001 as determined by One-way ANOVAs followed by Tukey’s post hoc test for multiple comparisons.

The observation that FBW7 associates with genes with VDREs in a 1,25D-dependent manner suggested that it may be recruited by the VDR to the E boxes of c-MYC target genes, whose expression is repressed by 1,25D. Our previous studies showed that 1,25D treatment over a 24h period largely abolished c-MYC binding to E boxes of target promoters and led to a dramatic increase in recruitment of MXD1 to the same sites, consistent with the opposing effects of hormone on c-MYC and MXD1 turnover and expression [16]. The same study also detected the VDR associated with the same promoters by ChIP [16]. Consistent with these findings, 6h treatment with 1,25D reduced the association of c-MYC with the *CDC25A* and *CDK4* promoters (Fig. 3A). Under the same conditions, association of tagged FBW7 with the same loci was detected by ChIP, with little apparent effect of 1,25D (Fig. 3B). ChIP assays also revealed an association of proteasome subunits SKP1 and RBX1 with the *CDC25A* and *CDK1* loci (Figs. S5B and C), and, similar to total FBW7 (Fig. 3B), there was a lack of apparent dependence on 1,25D treatment on SKP1 and RBX1 recruitment. However, such results likely represent a composite of dynamic, 1,25D-dependent interactions of FBW7 with changing levels of c-MYC, MXD1 and the VDR at these sites. To probe this further, reChIP assays were performed with an anti-HA antibody against FBW7. These experiments revealed an increased association of FBW7 with DNA-bound c-MYC after 6h of 1,25D treatment (Fig. 3C), even though DNA-bound c-MYC was in decline (Fig. 3A). Conversely, we observed a markedly decreased colocalization of FBW7 with MXD1 under the same conditions (Fig. 3C), as well as a decline in reChIP of the VDR with FBW7 associated with these sites (Fig. 3D). This is to be expected given that these sites are predominately occupied by MXD1 after extended 1,25D treatment [16], which would be refractory to FBW7 binding in the presence of 1,25D. ReChIP assays also revealed a 1,25D-dependent increase in ubiquitin associated with DNA-bound c-MYC (Fig. 3E), consistent with 1,25D inducing c-MYC turnover on DNA. Finally, depletion of FBW7 strongly reduced association of the VDR with the *CDC25A* E box (fig. S4B), consistent with its effects on binding of the receptor to VDRE-containing genes. These results are also highly consistent with the protein-protein interaction studies above describing the 1,25D-dependence of interactions of FBW7 with c-MYC, MXD1, and the VDR. They confirm that 1,25D differentially alters the association of c-MYC and MXD1 with E boxes, and reveal that hormone controls the recruitment of FBW7 and proteasomal subunits to these sites.

**Figure 3:**
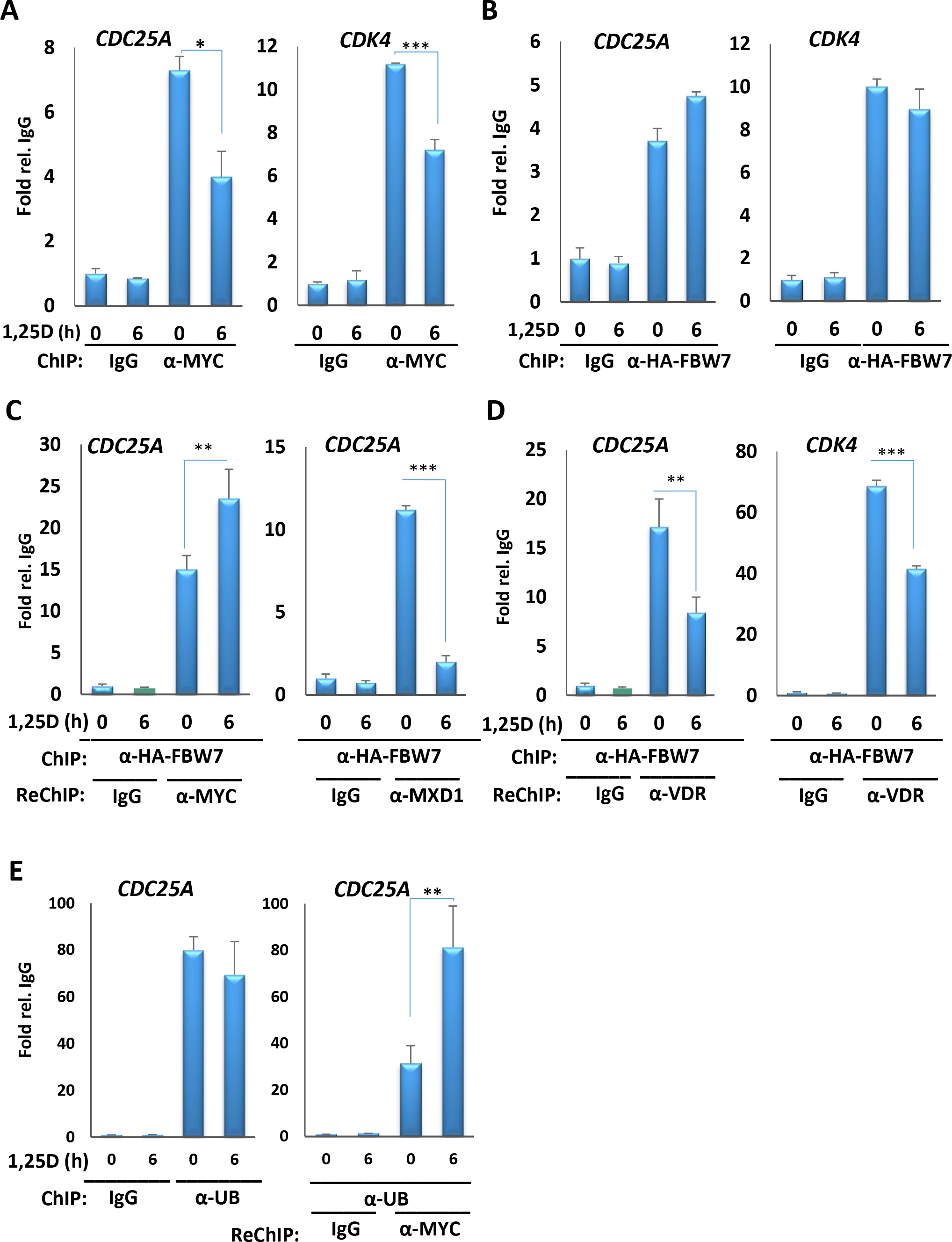
VDR and FBW7 are co-recruited to E-box motifs of c-MYC target genes. **A)** Analysis of c-MYC recruitment to E-boxes of target genes by ChIP-qPCR in SCC25 cells treated with 100nM 1,25D for 6 h. **B)** Analysis of HA-FBW7 recruitment to E-box motifs by ChIP-qPCR in HA-FBW7 SCC25 cells treated with 100nM 1,25D for 6 h. **C)** Analysis of the co-recruitment of HA-FBW7 and c-MYC or MXD1 to E-box motif of *CDC25A* by Re-ChIP in SCC25 cells after 6h 1,25D treatment. The first round of ChIP was for HA-FBW7, followed by second round of IP for c-MYC or MXD1. **D)** Analysis of the co-recruitment of HA-FBW7 and VDR to E-box motifs by re-ChIP assay in SCC25 cells. The first round of ChIP was for HA-FBW7, followed by second round of IP for VDR. **E)** Left: Analysis of the recruitment of ubiquitin by ChIP assay and corecruitment of ubiquitin and c-MYC by re-ChIP assay to E-box motif of *CDC25A* promoter after 6h 1,25D treatment in SCC25 cells. *P≤0.05, **P≤0.01, ***P≤0.001 as determined by One-way ANOVAs followed by Tukey’s post hoc test for multiple comparisons.

Results presented above show that the hormone-bound VDR enhances the association of FBW7 and c-MYC and blocks that with MXD1, and suggest that the VDR may alter the expression of other FBW7 target proteins. We examined the effect of 1,25D treatment on four well-characterized FBW7-regulated proteins, AIB1, Cyclin E, MCL1 and c-JUN [8], [9]. In all cases, 1,25D treatment of SCC25 cells strongly reduced protein expression (Fig. 4A). Complete loss of AIB1 expression was also observed in 1,25D-treated HACAT keratinocytes (Fig. 4B). Under the same conditions, 1,25D treatment only modestly affecting corresponding mRNA levels (Fig. 4C). Significantly, the 1,25D-dependent loss in AIB1 and Cyclin E expression in SCC25 cells was blocked by ablation of *FBW7* expression (Fig. 4D).

**Figure 4:**
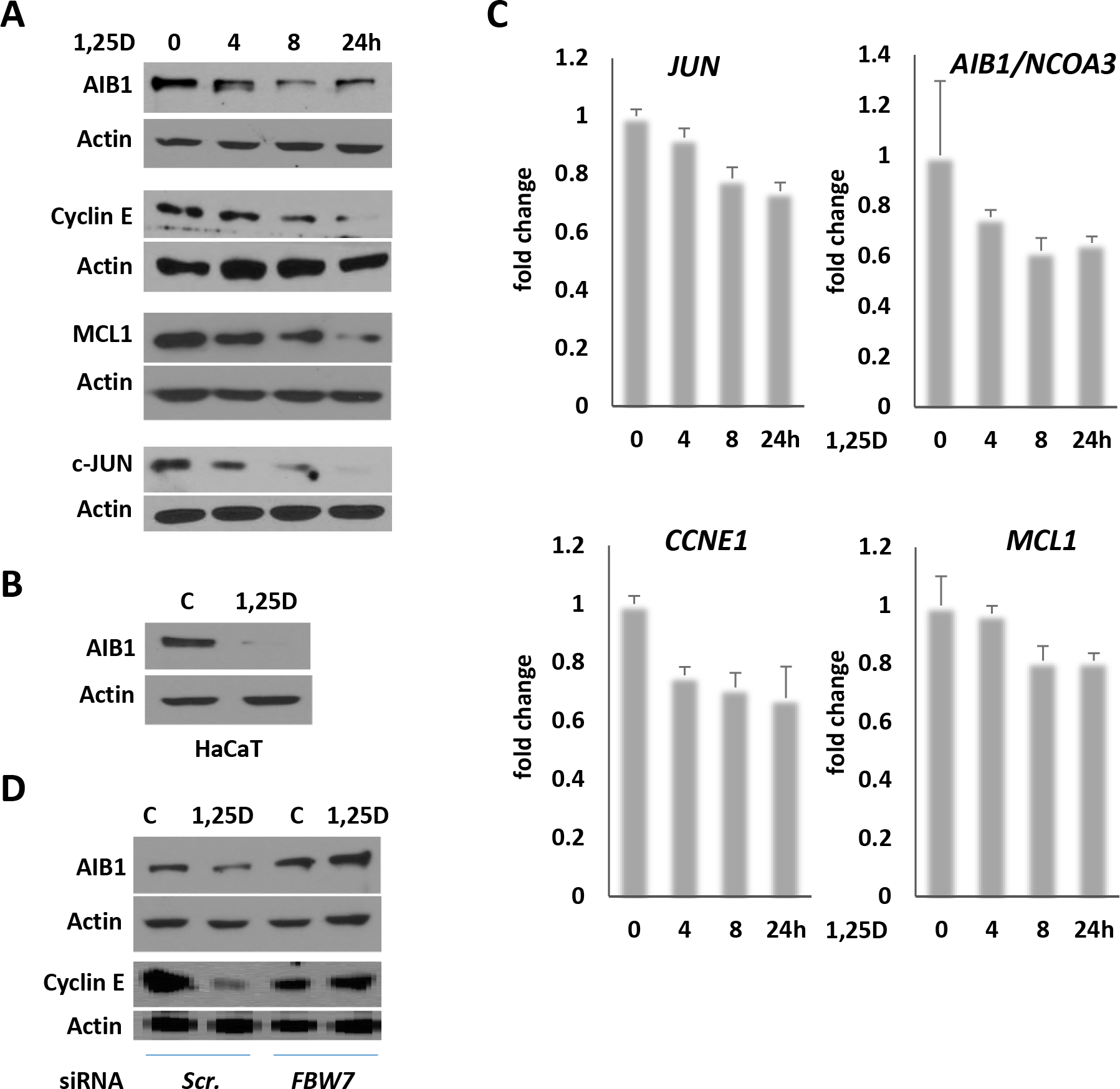
FBW7 target proteins controlling cell cycle progression are regulated by 1,25D. **A)** Western analysis of AIB1, Cyclin E, MCL1 and C-Jun levels in SCC25 cells following treatment with 100nM 1,25D. **B)** Western analysis of AIB1 in HaCaT cells following treatment with 100nM 1,25D for 24 h. **C)** RT-qPCR analysis of FBW7 target genes, *JUN*, *AIB1*, *CCNE1*, and *MCL1*, transcription in SCC25 cells following treatment with 100nM 1,25D. **D)** Western analysis of AIB1 and Cyclin E levels in SCC25 cells transfected with scrambled or *FBW7* siRNA and treated with vehicle or 100nM of 1,25D.

In conclusion, data presented above strongly suggest that the hormone-bound VDR is a key cofactor of FBW7, which is a tumor suppressor, and show that 1,25D regulates the association of FBW7 with two of its substrates, c-MYC and MXD1. The VDR interacts with FBW7 and c-MYC, and 1,25D enhanced the association of FBW7 with c-MYC as assessed by CoIP. Treatment with 1,25D also induced the association of FBW7 and ubiquitin with DNA-bound c-MYC as measured by re-ChIP assays. These results, summarized in Fig. 5, suggest that the hormone-bound VDR accelerates the proteasomal turnover of DNA-bound c-MYC, providing a mechanism for eliminating the protein and clearing E boxes for binding of MXD1, whose expression and DNA binding increase strongly in 1,25D-treated cells [16]. Other data presented provide evidence that the effects of 1,25D on FBW7 function as an E3 ligase are not limited to c-MYC and MXD1, as 1,25D treatment reduced protein, but not mRNA expression, of FBW7 targets AIB1, MCL1, cJUN and Cyclin E1, and ablation of FBW7 attenuated loss of AIB1 expression in 1,25D-treated cells. Consistent with the fact that many FBW7 substrates are drivers of cell cycle progression, ablation of FBW7 expression strongly compromised 1,25D-dependent cell cycle arrest. Taken together, these date strongly suggest that the hormone-bound VDR has widespread effects on FBW7-regulated proteasomal turnover, and that control of FBW7, which stimulates the degradation of several drivers of cell division, is a key component of the capacity of 1,25D to arrest cell proliferation.

**Figure 5:**
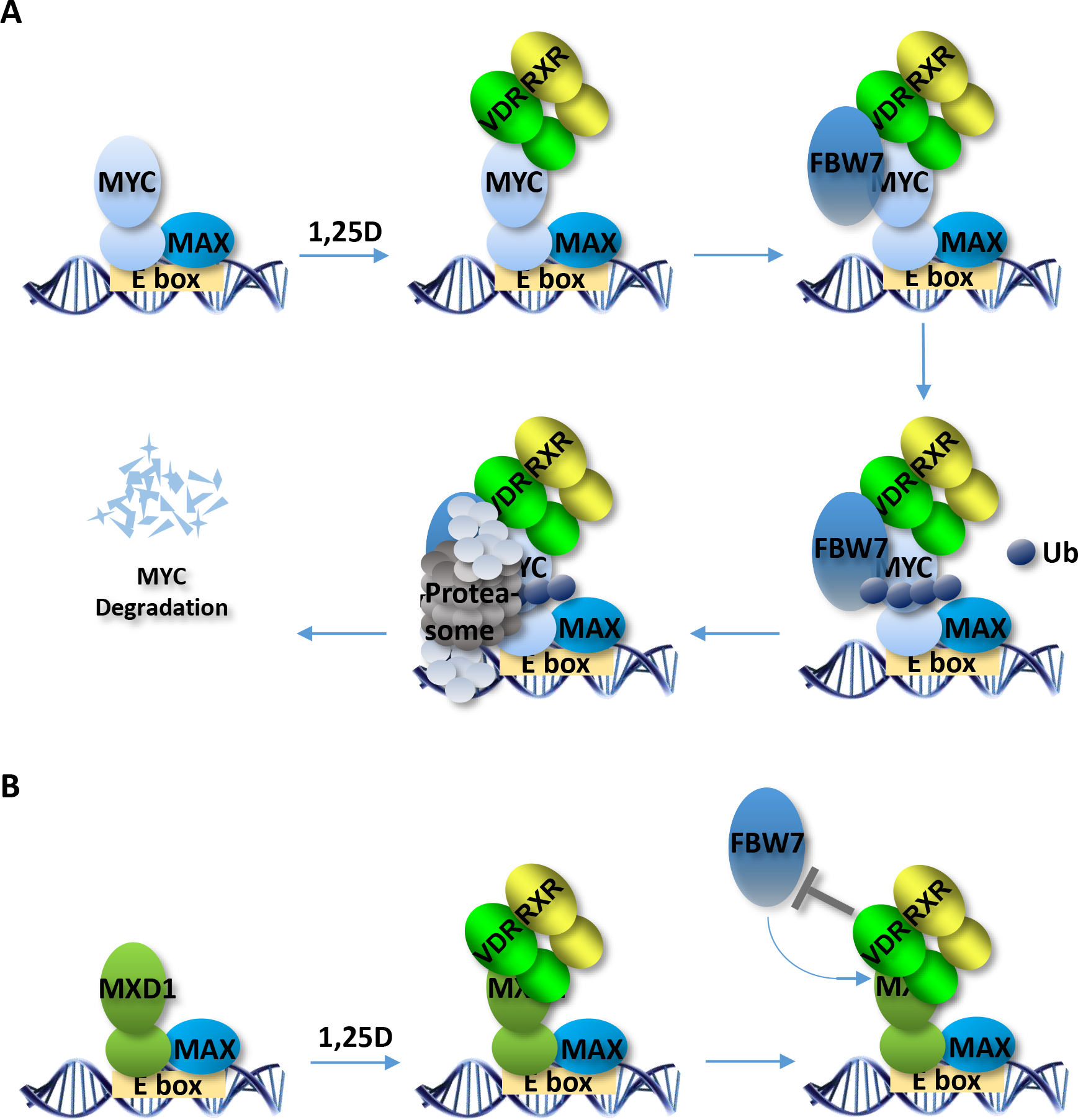
Schematic representation of the mechanism of recruitment and function of FBW7 E3 ligase complex on the promoter of VDR and MYC target genes.

These results are intriguing in light of epidemiological studies and preclinical data that link vitamin D deficiency to increased cancer risk [17]–[19], [24]–[30]. This includes, for example, a large prospective analysis linked vitamin D sufficiency to reduced total incidence and mortality from digestive cancers and leukemias [17]. Vitamin D deficiency represents a potentially widespread problem as, in the absence of adequate supplementation, most diets are vitamin D-poor. In addition, climate, extended vitamin D winters (inadequate solar UVB) in major population centers, conservative dress, and sun avoidance also contribute to vitamin D deficiency worldwide [31], [32]. However, while clinical and preclinical data point to cancer-preventive effects of vitamin D, much remains to be learned about the signaling events underlying its action. The findings that the hormone-bound VDR regulates substrate binding of FBW7 and accelerates the turnover of drivers of cell cycle progression provide key mechanistic support for the cancer preventive actions of vitamin D.

Sequence analysis revealed that the VDR has a motif corresponding to a consensus phospho-degron in an unstructured, VDR-specific insertion in the receptor ligand-binding domain. However, mutational analysis, CoIP experiments, and analysis of the effect of FBW7 depletion on VDR stability are not consistent with the VDR being a target of FBW7-regulated proteasomal turnover. It is not clear what the structure of the insertion region is, but the data suggest that the phospho-degron motif is sequestered and that FBW7 interacts with other site(s) on the VDR. The lack of stabilization of the VDR upon FBW7 depletion suggests that FBW7 associates with the VDR through region(s) independent of the substrate recognition domain. Unexpectedly, we also found that depletion of FBW7 reduces VDR gene and protein expression and receptor binding to cognate VDREs of target genes. In addition, FBW7 and the action of the proteasome were necessary for optimal VDR-dependent transactivation. We detected hormone-dependent recruitment of FBW7 and proteasome subunits to VDREs, and depletion of FBW7 or addition of proteasome inhibitors attenuated induction by 1,25D of multiple target genes.

Other studies have provided evidence for hormone-dependent recruitment of proteasome subunits to nuclear receptor target genes and their contribution to hormone-dependent transcriptional regulation. Estrogen receptor a (ERα) is degraded by the proteasome in a receptor-specific manner that is regulated by ligand binding, and proteasome inhibitors block estradiol-dependent transactivation [33],[34]. Subsequent work provided evidence that ERα-dependent transactivation and proteasome activity are intrinsically linked, and that proteasomal degradation of ERα occurs on estrogen-responsive promoters [35]. Analysis of androgen-regulated gene expression showed that recruitment of the MDM2 E3 ligase led to ubiquitination and degradation of the DNA-bound androgen receptor and that disruption of this process impaired hormone-dependent transactivation [36]. Similarly, proteasome inhibitors block transactivation by PPARγ, and receptor ubiquitination is enhanced by ligand binding [37]. Collectively, these studies provide strong evidence for a role of proteasomal degradation of nuclear receptors in the process of hormone-dependent transactivation, and suggest that receptor degradation may contribute to the putative turnover and cycling of transcription factors occurring during transactivation. However, it is unlikely that a similar FBW7-dependent mechanism of receptor turnover is operative on genes transactivated by the VDR, as FBW7 does not appear recognize the VDR as a typical substrate and does substantially regulate VDR stability. FBW7 may target other components of the transcription apparatus recruited by the VDR. If a similar mechanism of proteasomal degradation of the VDR occurs during transactivation it is likely mediated by other E3 ligase(s).

## Materials and methods

### Cell Culture

SCC25 cells were obtained from the American Type Culture Collection (ATCC) and cultured in DMEM/F12 (319-085-CL, Multicell) supplemented with 10% FBS. HaCaT cells (gift from Dr. Jean-JacquesLebrun’s lab) were cultured in DMEM (319-005-CL, Multicell) supplemented with 10% FBS. SCC25 cells with stable expression for tagged-FBW7 were generated using LeGO-iG2-Flag-FBW7, LeGO-iG2-HA-FBW7, and pLVX-IRES-Hyg-V5-FBW7 vectors. HEK 293-T17 (gift from Dr. Pelletier’s Lab) were cultured in DMEM (319-005CL, Multicell) supplemented with 10% FBS.

### Reagents

1α,25-Dihydroxyvitamin D3 (BML-DM200) was purchased from enzolifesciences, cycloheximide (C7698) were purchased from Sigma.

### Plasmids

In this study, the following constructs were generated: pcDNA 3.1-MXD1, pcDNA 3.1-MYC, pGEX-4T3-VDR, LeGO-iG2-Flag-FBW7, LeGO-iG2-HA-FBW7, and pLVX-IRES-Hyg-V5-FBW7.

### FBW7 knockdown

SCC25 cells were transfected with SMARTpool: ON-TARGETplus FBXW7 siRNA L-004264-00-0005 or scrambled siRNA (Dharmacon) for 24 h, using Pepmute transfection reagent (SL100566) from Signagen.

### RT-qPCR

Quantitative RT-PCR was performed with BrightGreen Express 2X qPCR MasterMix-ROX (Abmgood). Expression was normalized to the expression of Actin.

Primers for mRNA expression are as follows:

Actin F: CACCTTCACCGTTCCAGTTTT, R: AACCTAACTTGCGCAGAAAACAA;
MXD1 F:ACCTGAAGAGGCAGCTGGAGAA, MXD1 R:AGATAGTCCGTGCTCTCCACGT;
FBW7 F: CAGCAGTCACAGGCAAATGT, R: GCATCTCGAGAACCGCTAAC;
VDR F: CGCATCATTGCCATACTGCTGG, R: CCACCATCATTCACACGAACTGG;
CYP24A1 F: GCTTCTCCAGAAGAATGCAGGG, R: CAGACCTTGGTGTTGAGGCTCT;
CAMP F: GACACAGCAGTCACCAGAGGAT, R: TCACAACTGATGTCAAAGGAGCC;
SPP1 F: CGAGGTGATAGTGTGGTTTATGG, R: GCACCATTCAACTCCTCGCTTTC;
NOD2 F: GCACTGATGCTGGCAAAGAACG, R: CTTCAGTCCTTCTGCGAGAGAAC;
COX2 F: CGGTGAAACTCTGGCTAGACAG, R: GCAAACCGTAGATGCTCAGGGA;
AREG F: GCACCTGGAAGCAGTAACATGC, R: GGCAGCTATGGCTGCTAATGCA;
JUN F: CCTTGAAAGCTCAGAACTCGGAG, R: TGCTGCGTTAGCATGAGTTGGC;
AIB1 F: GGACTAAGCAACAGGTGTTTCAAG, R: ACTGGAGGACTTGAGCCAACAG;
CCNE1 F: TGTGTCCTGGATGTTGACTGCC, R: CTCTATGTCGCACCACTGATACC;
MCL1 F: CCAAGAAAGCTGCATCGAACCAT, R: CAGCACATTCCTGATGCCACCT;

### Immunoprecipitation and Western blot analysis

Cells were lysed with a lysis buffer (20 mM Tris, pH 7.5, 100 Mm NaCl, 0.5% Nonidet P-40, 0.5mM EDTA, 0.5 mM phenylmethylsulfonyl fluoride). 4 ng anti-c-Myc (D84C12, cell signaling) or MXD1(c-19, Santa Cruz) Antibodies were pre-bound for 2 hours to Dynabeads protein A, then was washed with lysis buffer and added to the lysate, followed by immunoprecipitation overnight. Beads were then washed five times with washing buffer (20 mM Tris, pH 7.5, 200 Mm NaCl, 1% Nonidet P-40, 0.5mM EDTA, 0.5 mM phenylmethylsulfonyl fluoride) and processed for Western blotting, performed with standard protocols. The following antibodies were used for western blot analysis: VDR (D-6), MXD1(c-19), HA (Y-11), C-Jun (H-79), Cyclin E (M-20) from Santa Cruz; AIB1 (5E11), E2F1 (3742), and c-MYC (D84C12) from cell signaling; Flag (M2) from sigma; Lamin-A (Gift from Dr. Stochaj’s lab).

### GST pull down assay

GST pull down assays were performed using the MagneGST Pull-Down System (Promega) according supplier’s instruction. BL21 bacteria were transformed with appropriate pGEX4T3 constructs and induced to express GST or GST fussed proteins. The total lysates of bacteria were used to pull-down in vitro translated proteins. The Pulled-Down proteins were analyzed by western blot assay.

### ChIP and Re-ChIP assays

ChIP assays were performed as described previously [16]. DNA fragments were purified with a PCR purification kit (Qiagen) and were analyzed by SsoFast-EvaGreen real-time PCR. Anti-HA (Y-11) from Santa Cruz, V5 (R960-25) from Thermo Fisher, MYC (9402), RBX1 (4397) and SKP1 (2156) from cell signaling and UB (FK2) from ENZO were used for ChIP. For re-ChIP assays, HA-FBW7 immunocomplexes were eluted by adding 40μl 10mM DTT for 30 min at 37°C. Supernatants were diluted 1:40 in dilution buffer (150 mM NaCl, 1% Triton X-100,2mM EDTA and 50 mM Tris-HCl, pH 8), and re-ChIPs were performed using anti-c-MYC (9402-Cell signaling), anti-MXD1(c-19-Santa Cruz) or anti- VDR (D-6-Santa Cruz) antibodies.

### Proliferation assays

Proliferation assay were performed by using Click-iT EdU Alexa Fluor 647 high-throughput imaging assay kit (Invitrogen) according to the manufacturer’s protocol. SCC25 cells were transfected with FBW7 or scrambled siRNAs and treated with 1,25D for 24 hours. Cells were incubated with Alexa-EdU for 1 hour and fixed with 3.7% formaldehyde. DAPI was used to stain the nucleus. Images were captured by High-content screening (HCS) microscope using ImageXpressMicro program and analyzed with MetaXpress.

### Statistical Analysis

All experiments are representative of 3-5 biological replicates. Unless otherwise indicated in the figures, statistical analysis was conducted using the program SYSTAT13 by performing one-way analysis of variance (ANOVA) followed by the Tukey test for multiple comparisons as indicated: *P≤0.05, **P≤0.01, ***P≤0.001.

## Acknowledgments

This work was supported by an operating grant from the Cancer Research Society to J.H.W. R.S-T. and B.M. were supported by McGill University internal studentships. We are grateful to Dr. Natacha Rochel, IGBMC, Illkirch, France for the graphic of the DNA-bound VDR/RXR.

## Author Contributions

R.S.-T. performed experiments, contributed to experimental design, and wrote part of the manuscript. B.M. performed ChIP, reChIP assays. H.W. helped in performing some of the experiments. N.R. produced the 3-dimensional model of showing the position of the phospho-degron motif in the VDR. J.H.W. contributed to experimental design and wrote and edited the manuscript.

## Supplementary Figures

**Figure S1:**
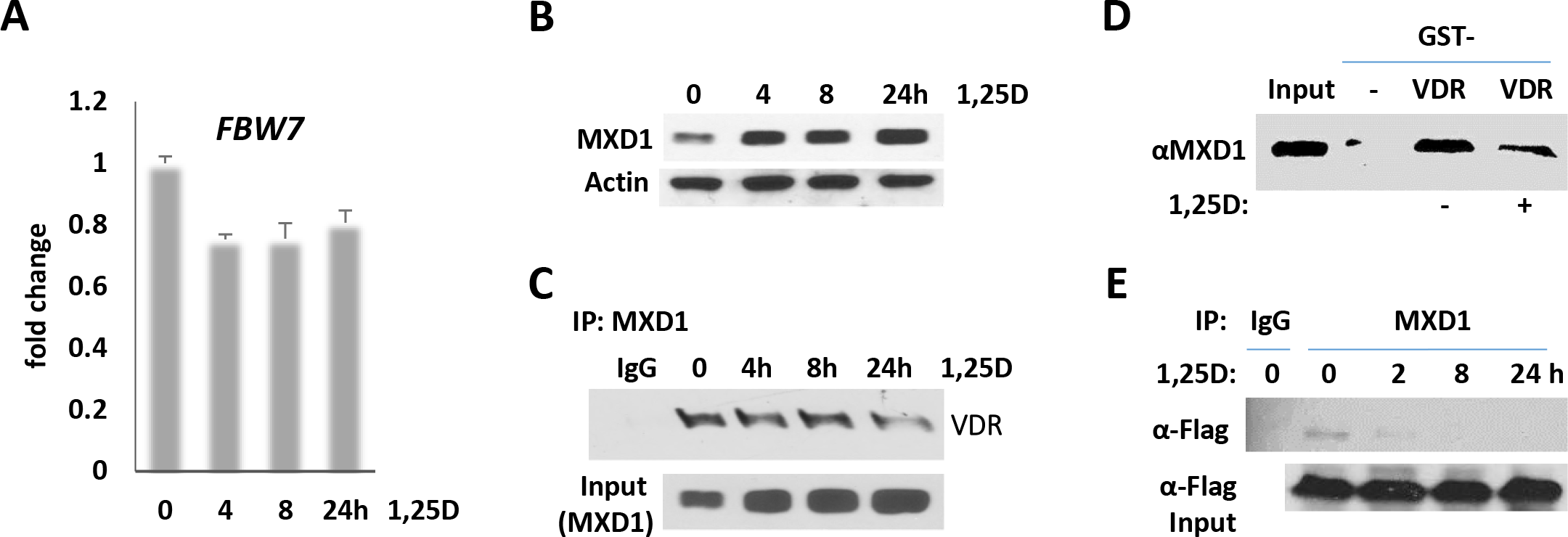
**A)** RT-qPCR analysis of *FBW7* gene transcription in SCC25 cells following treatment with 100nM of 1,25D for 4, 8, and 24 hours. **B)** Western blot analysis of MXD1 levels in SCC25 cells following treatment with 100nM of 1,25D for 4, 8, and 24 hours. **C)** Western blot analysis of VDR Co-IPed with IgG or MXD1, following treatment of SCC25 cells with 100nM of 1,25D for 4, 8, and 24 hours. **D)** Western blot analysis of MXD1 pulled down by GST or GST-VDR in the presence of vehicle or 1,25D. BL21 bacteria were transformed with pGEX4T3 or pGEX-4T3-*VDR* and induced to express GST or GST fussed VDR. MXD1 protein was translated *in vitro* from *pCDNA*3.1-*MXD1*. The bacterial lysates were used to pull down the in vitro translated MXD1. **E)** Western blot analysis of Flag-FBW7 Co-IPed with IgG or MXD1, following treatment of SCC25 cells with 1,25D. SCC25 cells were transfected with empty vector or Flage-FBW7 and treated with 100nM 1,25D. MXD1 was IPed from SCC25 cells lysate. The total lysate as “input” and IPes were probed for Flag.

**Figure S2:**
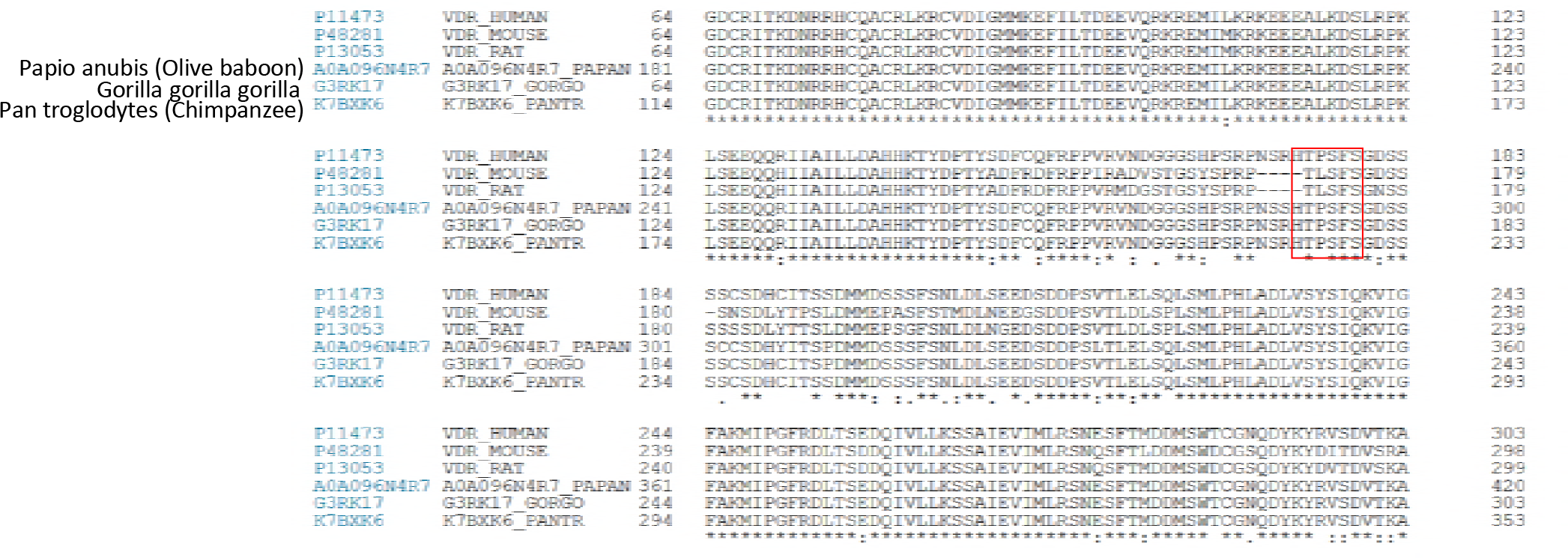
Alignment of phospho-degron DNA motif in VDR genes from different species.

**Figure S3:**
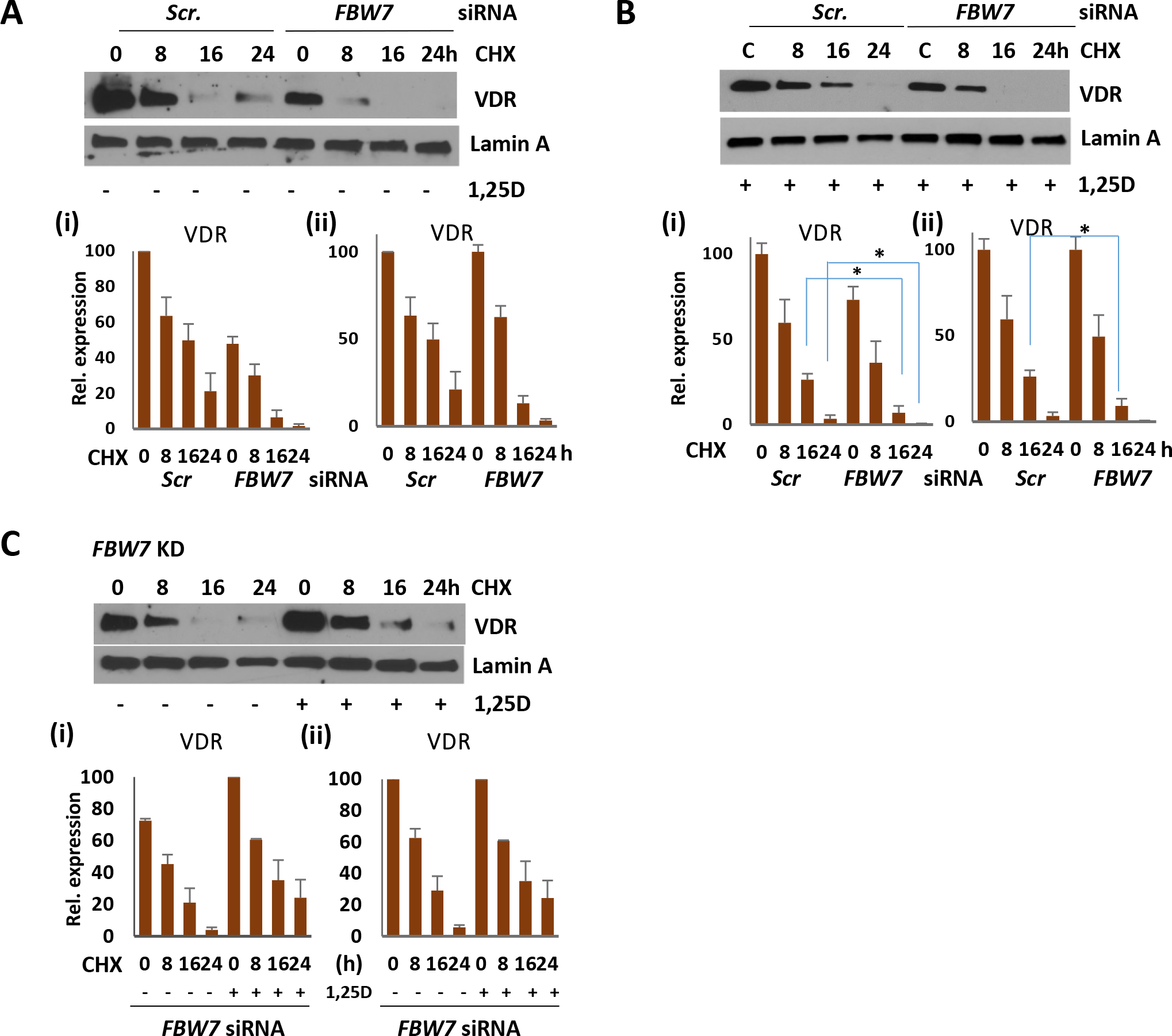
**A, B)** Top: Western analysis of VDR in protein stability assay. 4 μg/ml of Cycloheximide (CHX) were added for 8, 16 or 24h. Bottom: Quantification of Western analyses of 3 independent experiments. (i) VDR levels were normalized to those of scrambled siRNA transfected cells at time 0 (ii) VDR levels were normalized to those at time 0 in both Scrambled and *FBW7* siRNA transfected cells. **A)** SCC25 cells were transfected for *FBW7* or scrambled siRNA. Lamin-A was used as an internal control. **B)** SCC25 cells were transfected for *FBW7* or scrambled siRNA and treated with 100nM 1,25D. **C)** Analysis of the effect of 1,25D on VDR stability in SCC25 cells depleted of *FBW7*. Top: Western analysis of VDR in protein stability assay. 4 μg/ml of Cycloheximide (CHX) were added for 8, 16 or 24h. Bottom: Quantification of Western analyses of 3 independent experiments. (i) VDR levels were normalized to those of vehicle treated cells at time 0 (ii) VDR levels were normalized to those at time 0 in both vehicle and 1,25D treated cells. *P≤0.05 as determined by One-way ANOVAs followed by Tukey’s post hoc test for multiple comparisons.

**Figure S4:**
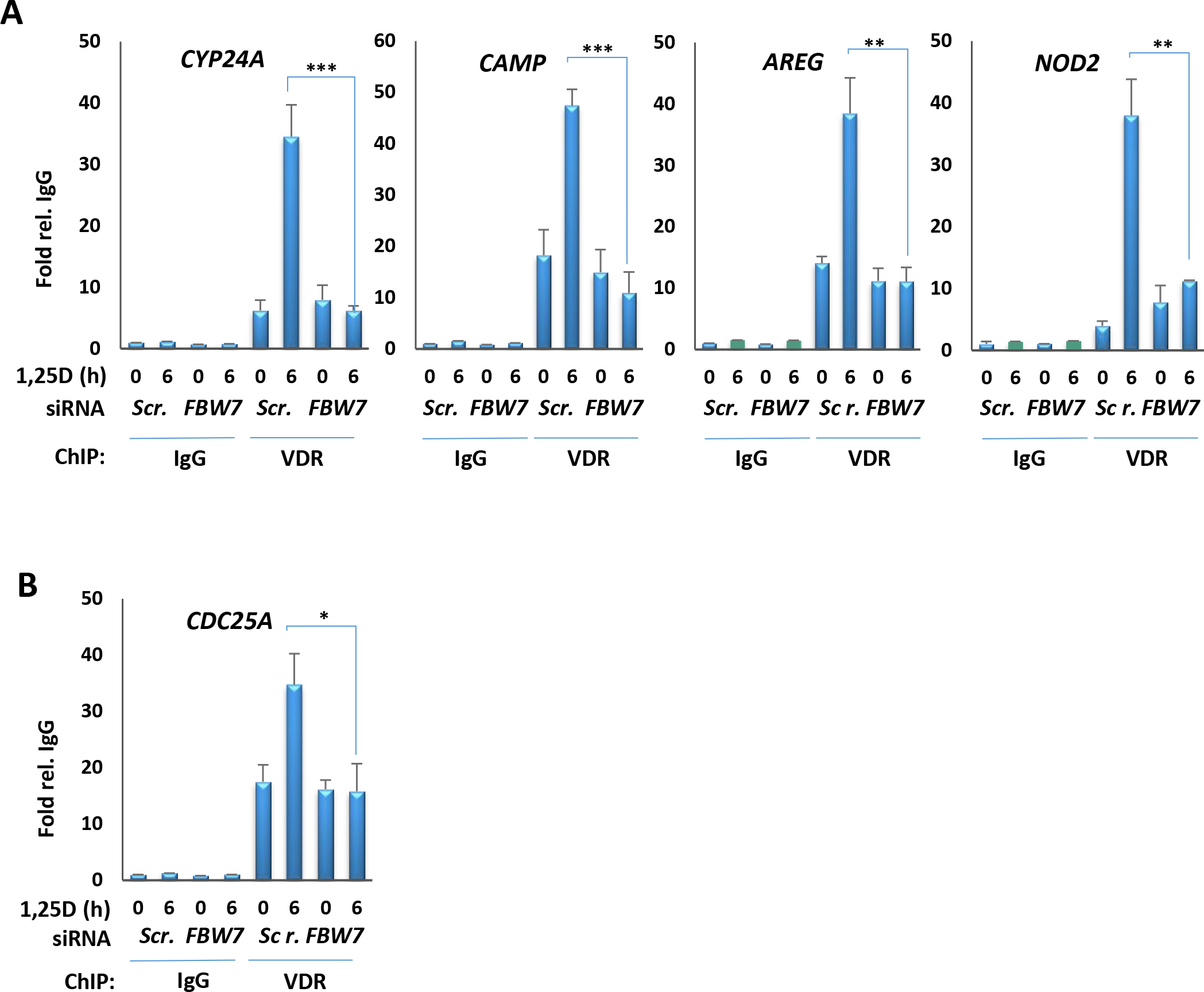
**A)** ChIP analysis of VDR binding to VDREs in control and FBW7-depleted cells. **B)** ChIP analysis of VDR association with the E-box of *CDC25A* in control and FBW7-depleted cells. *P≤0.05, **P≤0.01, ***P≤0.001 as determined by One-way ANOVAs followed by Tukey’s post hoc test for multiple comparisons.

**Figure S5:**
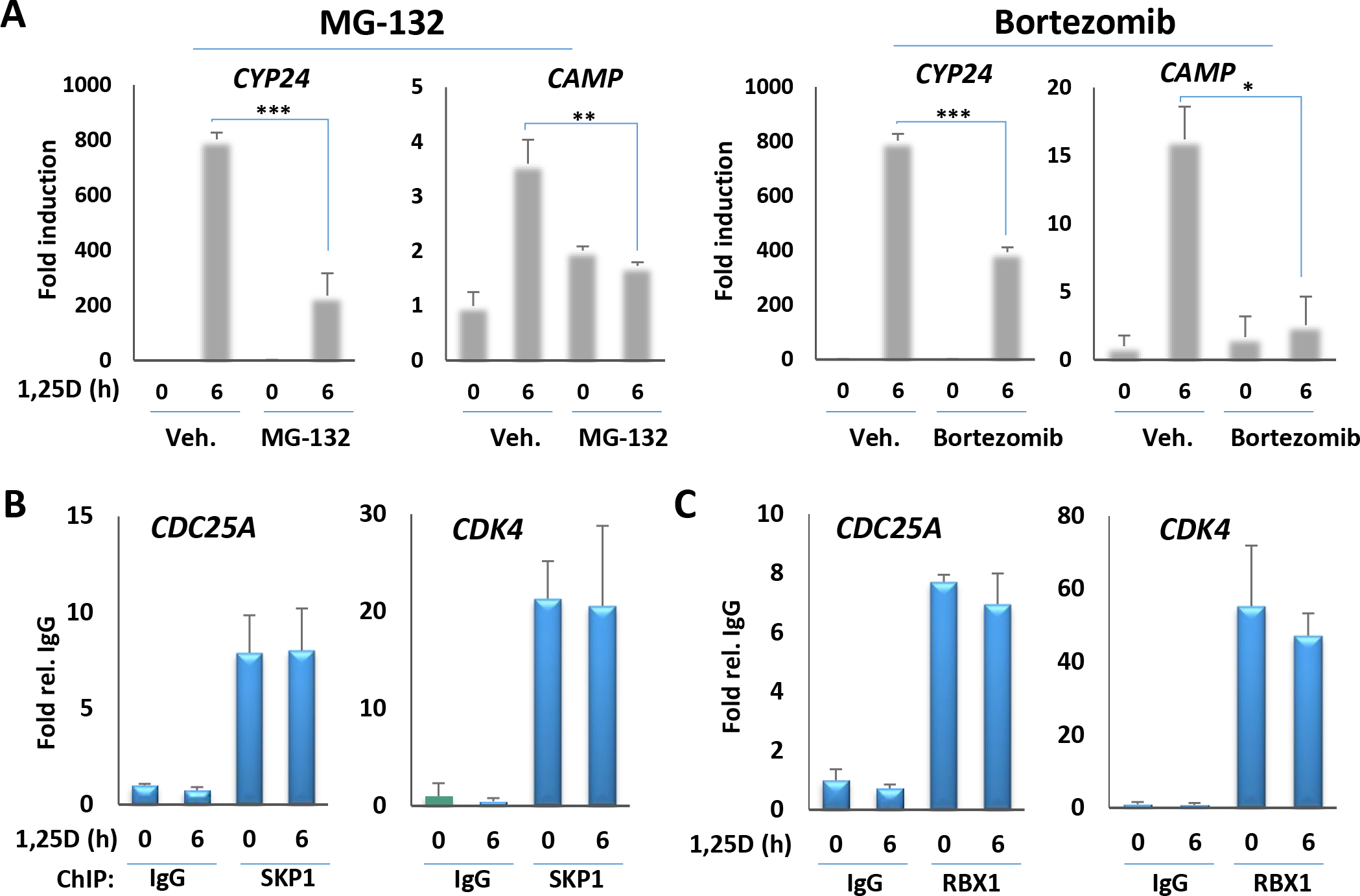
**A)** RT-qPCR analysis of VDR target genes transcription in SCC25 cells in the absence or presence of proteasome inhibitor MG-132 or Bortezomib. **B)** Analysis of recruitment of FBW7 E3 ligase complex component SKP1 to E-boxes of MYC target genes, *CDC25A* and *CDK4*, by ChIP/qPCR in SCC25 cells. **C)** Analysis of recruitment of FBW7 E3 ligase complex component RBX1 to E-boxes of MYC target genes, *CDC25A* and *CDK4*, by ChIP/qPCR in SCC25 cells. *P≤0.05, **P≤0.01, ***P≤0.001 as determined by One-way ANOVAs followed by Tukey’s post hoc test for multiple comparisons.

